# Tight association between microbial eukaryote and *Imitervirales* communities in the Pacific Arctic Ocean

**DOI:** 10.1101/2021.09.02.458798

**Authors:** Jun Xia, Sohiko Kameyama, Florian Prodinger, Takashi Yoshida, Kyoung-Ho Cho, Jinyoung Jung, Sung-Ho Kang, Eun-Jin Yang, Hiroyuki Ogata, Hisashi Endo

**Affiliations:** Institute for Chemical Research, Kyoto University, Gokasho, Uji 611-0011, Japan; Faculty of Environmental Earth Science, Hokkaido University, N10W5 Sapporo, Hokkaido 060-0810, Japan; Graduate School of Agriculture, Kyoto University, Kitashirakawa-Oiwake, Sakyo-ku, Kyoto 606-8502, Japan; Korea Polar Research Institute, Incheon 21990, Republic of Korea

## Abstract

Viruses are important regulatory factors of marine microbial community including microeukaryotes. However, little is known about their role in the northern Chukchi Sea of the Arctic basin, which remains oligotrophic conditions in summer. To elucidate linkages of microbial eukaryotic community with viruses as well as environmental variables, we investigated the community structures of microeukaryotes (3–144 µm and 0.2–3 µm size fractions) and *Imitervirales* (0.2–3 µm size fraction), a major group of viruses infecting marine microeukaryotes. Surface water samples were collected at 21 ocean stations located in the northeastern Chukchi Sea (NECS), an adjacent area outside the Beaufort Gyre (Adjacent Sea; AS), and two melt ponds on sea ice in the summer of 2018. At the ocean stations, nutrient concentrations were low in most of the locations expect at the shelf in the AS. The community variations were significantly correlated between eukaryotes and *Imitervirales*, even within the NECS characterized by relatively homogeneous environmental conditions. The association of the eukaryotic community with the viral community was stronger than that with geographical and physicochemical environmental factors. These results suggest that *Imitervirales* are actively infecting their hosts even in cold and oligotrophic sea water in the Arctic Ocean.

## Introduction

The Arctic Ocean is the smallest, shallowest, and coldest ocean on earth. It covers several seas including the Chukchi Sea and the Beaufort Sea. Sea ice cover the central area of the ocean slightly in summer and almost completely in winter, due to the extreme seasonality in the receipt of solar radiation in the polar area. Anthropogenic climate change is now accelerating and has a strong impact on the Arctic Ocean (Lannuzel et al., 2020; Stroeve et al., 2007). Due to the ice-albedo feedback mechanism, sea ice retreat to a greater extent in summer than before (Curry et al., 1995; Kashiwase et al., 2017; Lindsay et al., 2012). The highest surface air temperature during the past 120 years was recorded for the period between 2014 and 2019 (Perovich & Jones, 2019). Upper sea water was also freshened as a result of accelerating sea ice melting, and this situation is predicted to continue (Kwok & Cunningham, 2010; Münchow, 2016; Shu et al., 2018). In such changing environmental conditions, how microorganisms form and alter their community structures through intricate interactions between them and with surrounding environments is an important general issue.

Microbial eukaryotes play a fundamental role in the marine ecosystem (de Vargas et al., 2015; Worden et al., 2015). Being integrated in the food web, they drive biogeochemical cycles by contributing to primary production (Falkowski et al., 1998; Field et al., 1998) and transferring fixed carbon to the higher trophic levels (Sherr et al., 2007). Primary production in surface seawater of the Arctic Ocean estimated by satellite remote sensing significantly increased from 1998 to 2018, presumably due to the climate change-induced environmental modifications such as sea ice loss and an increase of nutrient influx (Lewis et al., 2020). However, another study indicated that the abundance of nanophytoplankton (2–20 µm) decreased from 2004 to 2008 in the Canada Basin while that of picophytoplankton (i.e., cell size of 0.2–2 µm) increased because of the decrease in nutrient concentrations (Li et al., 2009).

A first molecular biological characterization of the microbial eukaryotic community in the Arctic Ocean has been reported fifteen years ago (Lovejoy et al., 2006). Since then, several groups have carried out molecular barcoding studies to investigate the microbial eukaryotic community in a variety of areas of the Arctic Ocean such as the Chukchi Sea, the Beaufort Sea, West Spitsbergen, the Amundsen Gulf, the Bering Strait, Greenland and the central Arctic Ocean (Comeau et al., 2011; Kilias et al., 2014; Marquardt et al., 2016; Monier et al., 2013; Onda et al., 2017; Terrado et al., 2009; Zhang et al., 2015). Strong seasonality has been revealed through the annual data of 18S rDNA derived from arctic surface water samples, with dominant microbial eukaryotic groups being largely different between summer and winter (Marquardt et al., 2016). Composition of the microbial eukaryotic community in the Arctic Ocean is also shown to vary across water masses and environments with different physicochemical parameters and nutrient concentrations (Hamilton et al., 2008; Joli et al., 2018; Kilias et al., 2014; Thaler & Lovejoy, 2013). These results collectively suggest the importance of environmental conditions in constructing microeukaryotes at large time and spatial scales.

Apart from the physicochemical properties, viruses are thought to be a key effector of the communities of their microbial hosts in marine ecosystems (Middelboe & Brussaard, 2017; Rohwer & Thurber, 2009; Suttle, 2007). *Imitervirales*, belonging to the phylum *Nucleocytoviricota* (also known as nucleocytoplasmic large DNA viruses, NCLDVs) (Iyer et al., 2006), is one of the most dominant orders of viruses infecting diverse microbial eukaryotes in the ocean (Endo et al., 2020). A previous study showed a tight association between the community of NCLDVs and that of microbial eukaryotes based on a global metagenomic dataset (Endo et al., 2020). However, as the result was based on a global scale dataset, the observed association was expected from substantial differences in the host eukaryotes inhabiting geographically distant and environmentally distinct locations. We consider that investigating such virus-host associations at a smaller geographic and time scales would provide further insights into the possible regulatory role of viruses on the host community structure. However, such local studies are currently scarce for *Imitervirales* (or NCLDVs) (Clerissi et al., 2012; Sandaa et al., 2018), and to our knowledge, there is no study investigating both *Imitervirales* and eukaryotic communities in the Arctic Ocean. We reasoned that examining whether the two communities are associated with each other or not in geographically close locations with similar environmental conditions and in the same period would lead to a better understanding of the interactions between *Imitervirales* and microbial eukaryotes. If virions persistently remain in an environment for a long period of time, then we would not expect a strong association between *Imitervirales* and microbial eukaryotes. In contrast, if viruses actively infect various eukaryotes, then a strong eukaryote-viral association would emerge.

In this study, we conducted a high spatial resolution sampling of microbial DNA from the surface water during the 2018 cruise of the Korean ice breaking research vessel (IBRV) Araon. We investigated the community structures of microeukaryotes and *Imitervirales* in the basin region of the Chukchi Sea (the northeastern Chukchi Sea; hereafter NECS) as well as in an adjacent sea (AS) outside the Beaufort Gyre and melt ponds. The surface water of the NECS is characterized by the low salinity and nutrients under the influence of the Beaufort Gyre system, making it distinct from the AS areas. To gain insight into the interdependence of the eukaryotic and *Imitervirales* communities, we analyzed the statistical relationships between the eukaryotic and *Imitervirales* communities while controlling the effects of environmental and geographical characteristics in the two different environmental regimes (the “stable” NECS and the “dynamic” AS).

## Results

### Water characteristics and environmental factors

Twenty-one oceanic sampling stations (surface seawater samples) were classified into two groups depending on the geographical locations and the temperature-salinity (TS) diagram: the NECS (in the regions of Chukchi Plateau and Canada basin) and the AS (Figs 1 and S2).

**Fig. 1.**
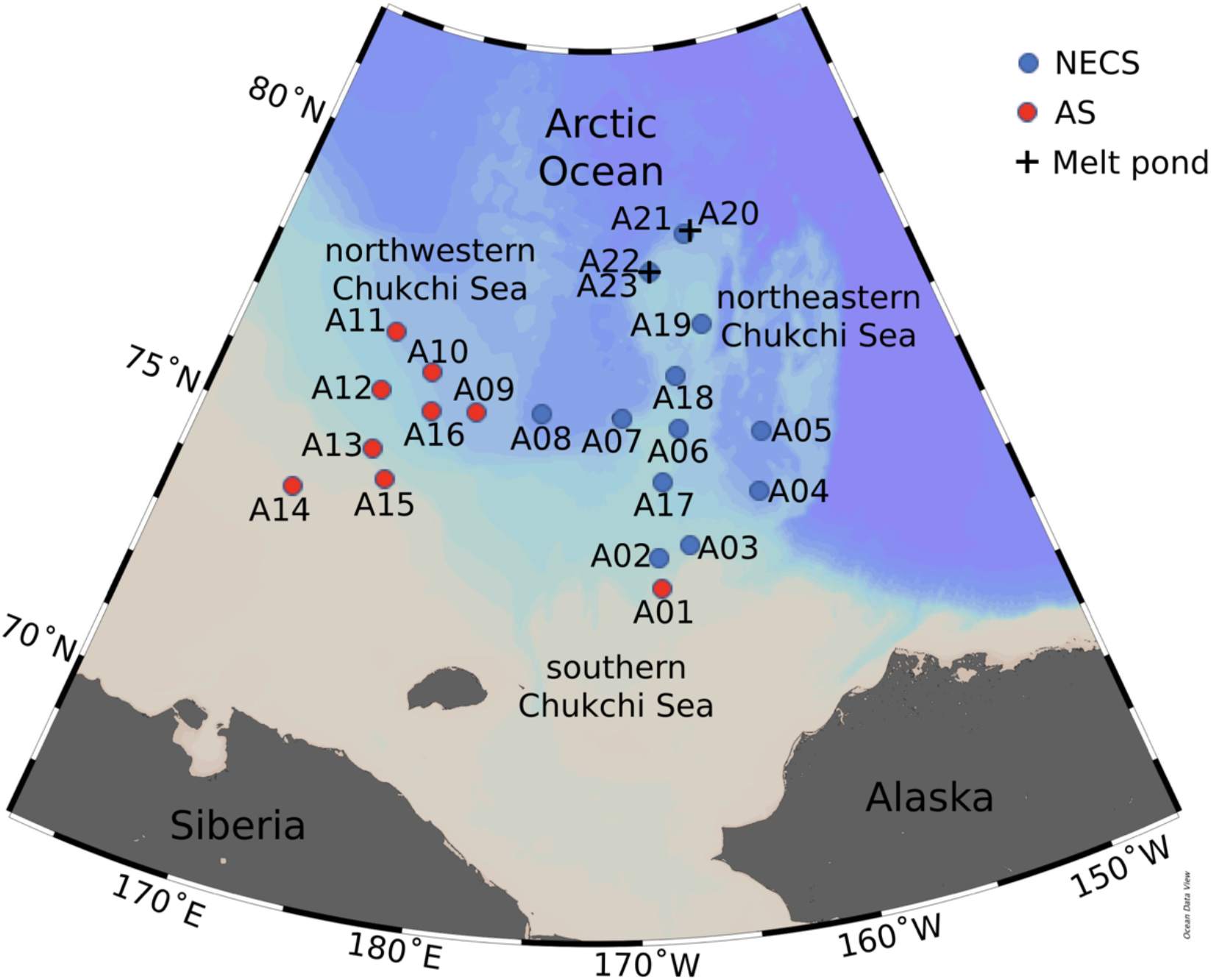
Map of Arctic sampling stations of this study. Symbol colors represent water types with different characteristics influenced by current system in this area. NECS: northeastern Chukchi Sea; AS: adjacent sea outside Beaufort Gyre.

Of the measured physical parameters (Supplement Table S1), temperature showed little difference among stations, but salinity showed relatively large differences (Supplement Table S1, Fig. 2A and B). Sea surface temperatures (SST) in the NECS (−0.99°C on average) was slightly higher than those in the AS (−1.11°C on average), and the salinity in the NECS (27.98 psu on average) was substantially lower than the AS (30.05 psu on average). Sea ice coverage in each station was 0% to 89.5% (Supplement Table S1).

**Fig. 2.**
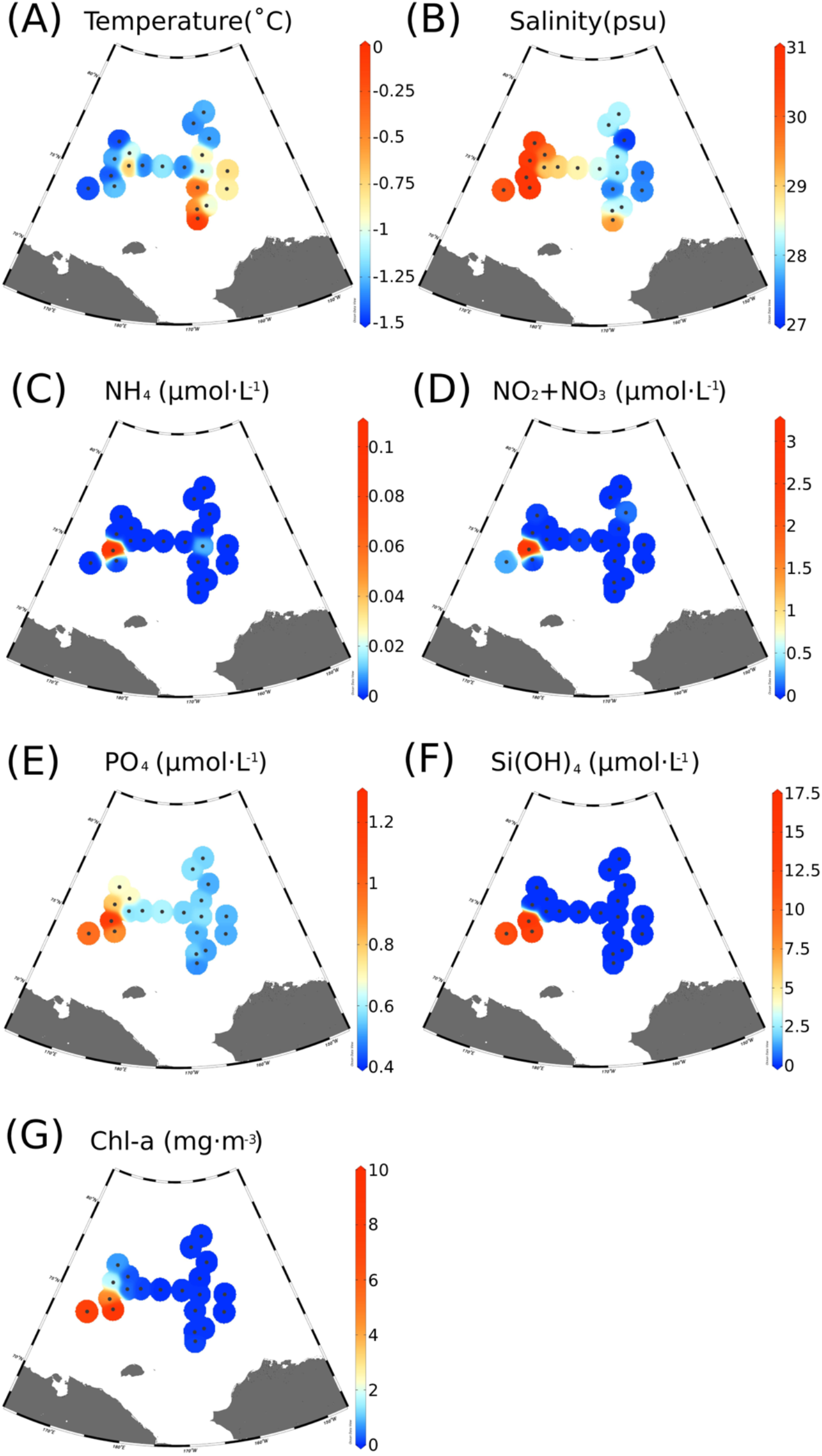
Physical and chemical environmental variables among sampling sites: (A) temperature (°C); (B) salinity (psu); (C) NH_4_ (µmol·L^-1^); (D) NO_2_ and NO_3_ (µmol·L^-1^); (E) PO_4_ (µmol·L^-1^); (F) Si(OH)_4_ (µmol·L^-1^); (G) Chl *a* (mg·m^-3^).

Concentrations of nutrients (ammonia nitrogen, nitrate+nitrite, phosphate, and silicate) as well as Chl *a* were measured for each location. Most of the sampled area was oligotrophic, while water conditions of three “bloom” sites (stations A13, A14, A15) in the AS presented high nutrient concentrations. The nutrient and Chl *a* concentrations for the bloom stations (on average: nitrate + nitrite: 1.17 µmol·L^-1^; phosphate: 1.02 µmol·L^-1^; silicate: 14.15 µmol·L^-1^; chlorophyll *a*: 7.23 mg·m^-3^) were much higher than those in other stations (on average: nitrate+nitrite: 0.14 µmol·L^-1^; phosphate: 0.52 µmol·L^-1^; silicate: 0.01 µmol·L^-1^; chlorophyll *a*: 0.17 mg·m^-3^). Ammonia concentration was relatively high at the station A13 (0.11 µmol·L^−1^), while it was less than 0.02 µmol·L^−1^ in other stations.

### Amplicon sequences

Sequence information and the number of ASVs were summarized in Supplement Table S2. The number of ASVs from each sample before subsampling is provided in Fig. S1. For the 3-144 µm eukaryotic community, 45,588 to 214,775 reads obtained from individual samples were subsampled at the minimum number of reads across different samples (i.e., 45,588 reads), and then grouped into 107 to 390 eukaryotic 18S non-singleton ASVs with the mean proportion of raw read usage being 37%. For the 0.2-3 µm eukaryotic community, subsampling was performed at 72,359 reads, which was grouped into 106 to 385 eukaryotic 18S ASVs. For the *Imitervirales* community, subsampling was also performed at 26,638 reads, which generated 244 to 525 *Imitervirales polB* ASVs per sample.

### Composition and local diversity of eukaryotic and *Imitervirales* communities

We first investigated eukaryotic communities by excluding sequences from metazoa and fungi, because they have different lifestyles and ecological functions from protists. The community compositions were different between large (3-144 µm) and small (0.2-3 µm) size-fractions (Fig. 3A and B). Eukaryotic communities of the large size fraction were dominated by dinoflagellates (36.6% on average), diatoms (11.4%) and other marine alveolates (29.7%, include ciliates and protaveolata), while those of small size fraction were dominated by ciliates (28.5%), chlorophytes (19.8%) and picozoa (10.8%). In the large size fractions, lower proportion of dinoflagellates occurred in the AS sites than in the NECS sites (ANOVA followed by Tukey post hoc tests, *p*<0.01), especially in the three bloom samples (ANOVA followed by Tukey post hoc tests, *p*<0.05). On the contrary, diatoms tended to be more abundant in the AS sites than in the NECS sites (ANOVA followed by Tukey post hoc tests, *p*<0.01). In the small size fraction, although the dominant ciliates had little difference among all the samples, chlorophytes showed higher proportion in the AS bloom samples and the NECS samples than in other samples of the AS sites (ANOVA followed by Tukey post hoc tests, *p*<0.01). Another unique feature of the bloom sites was that there was almost no picozoa sequences in these samples, while the picozoa represented one of the abundant phyla in the other samples.

**Fig. 3.**
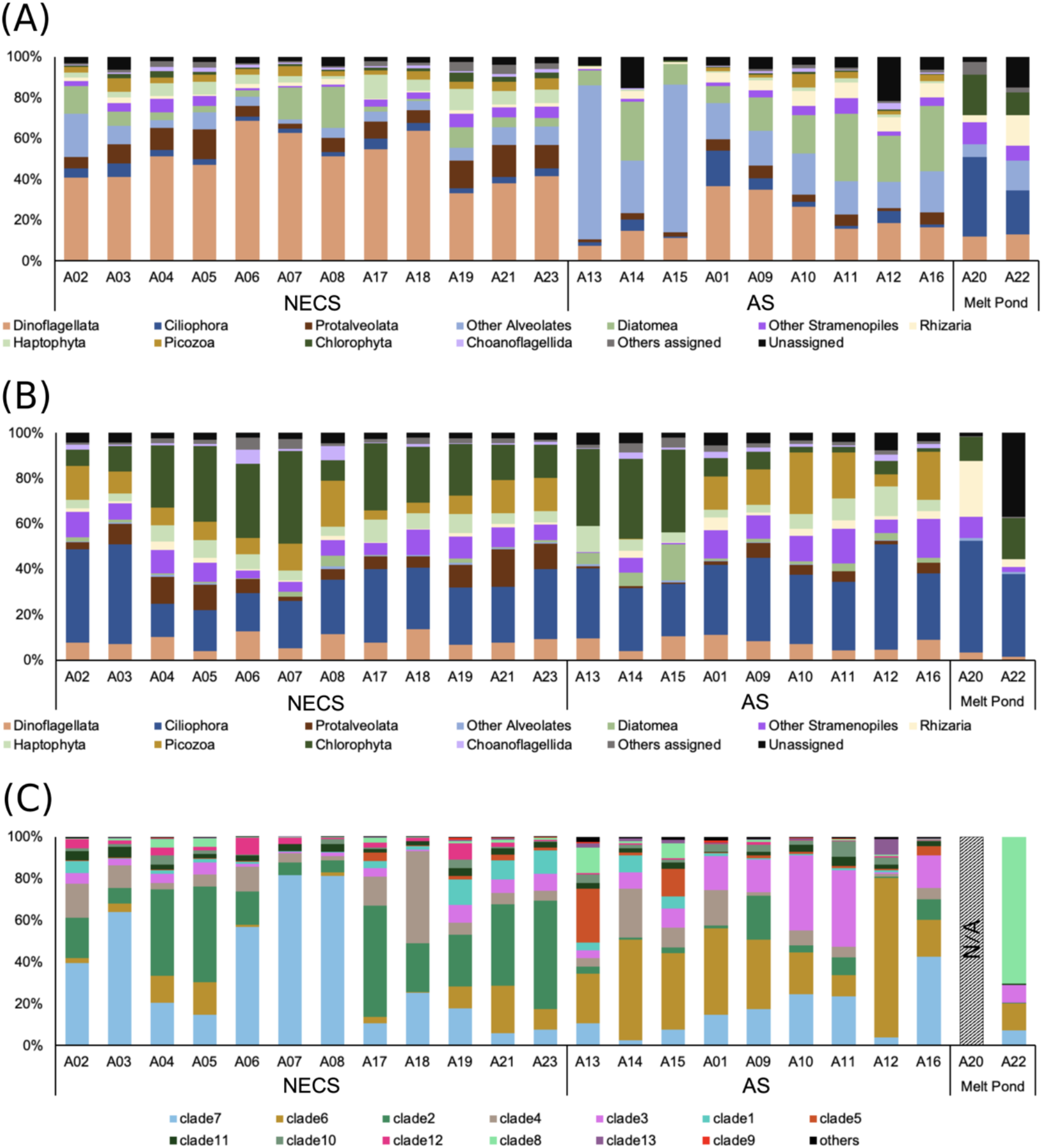
Community compositions of eukaryotes and *Imitervirales*. Relative compositions of eukaryotes at phylum level in (A) 3-144 µm fraction and (B) 0.2-3 µm fraction and (C) *Imitervirales* in the clade level. The color of each clade of the *Imitervirales* is the same with phylogenetic tree (Fig. S5). Fungi and Metazoa sequences were removed from eukaryotic sequences.

As for metazoa and fungal communities (Fig. S3), copepods were the most dominant (20.0% on average) in the 3-144 µm size fraction samples. As for protist community, the most abundant ASVs (>10% in at least one sample) in the large size fraction belonged to *Heterocapsa* (dinoflagellate), *Chytriodinium* (dinoflagellate), *Gyrodinium* (dinoflagellate), while those in the small size fraction were *Micromonas* (chlorophyte), *Oligotrichia* (ciliate), *Chytriodinium* (dinoflagellate), *Phaeocystis* (haptophyte), *Chaetoceros* (diatom) and *Carteria* (chlorophyte).

*Imitervirales* ASVs were mapped onto a larger set of *polB* sequences from the *Tara* Oceans dataset and classified into 13 clades (Fig. S5). Clades 7 (28.6%) was the most abundant clade, followed by clade 6 (20.2%) and clade 2 (17.0%) (Fig. 3C). In the bloom sites, particularly high relative abundances were shown for clade 5 and 6. In the other AS sites, clade 3 which includes the OLPVs (Organic Lake Phycodnavirus 1 and 2) showed higher proportions than in the NECS. In the NECS samples, clade 2 showed high proportions (28.0% on average). Clear difference was found between the communities in the two aquatic habitats (sea water and melt pond water) for the eukaryotic and *Imitervirales* communities (Fig. 3A–C and S6). *Imitervirales* communities in the Arctic Ocean were clearly distinguished with those obtained from subtropical coastal sea water and hotspring samples by the same amplicon method (Fig. S6) (Li et al., 2019; Prodinger et al., 2020; Prodinger et al., unpublished). In the samples of the present study, *Imitervirales* communities were classified into three groups: Arctic seawater, Arctic algae bloom related seawater and melt pond water (Fig. S6). The sites in the NECS and AS shared 702 common *Imitervirales* ASVs, while 357 and 871 unique ASVs were detected in the NECS and AS sites, respectively (Fig. S7A). It was also shown that 515 *Imitervirales* ASVs were shared between the bloom sites (A13, A14 and A15) and non-bloom sites (Fig. S7B), while 319 and 1,096 ASVs were unique to the bloom sites and non-bloom sites, respectively.

Shannon’s diversity index was calculated for each community (Fig. S4). Diversity of eukaryotic communities in the large size fraction showed the same variation trend as those in the small fraction among different samples. The three bloom sites in the AS had statistically lower diversity than others in both the large (ANOVA followed by Tukey post hoc tests, *p*<0.01) and small eukaryotic communities (ANOVA followed by Tukey post hoc tests, *p*<0.01). The bloom sites had higher diversity of *Imitervirales* (4.34) than other sites (3.83) on average, although it was not statistically significant (ANOVA, *p*=0.068) (Fig. S4D).

### Correlations with eukaryotic 18S community

Result of dbRDA (Fig. 4) and Spearman’s rank correlation (Supplement Table S5) demonstrated that salinity and longitude were the two most significant variables in predicting the Chl *a* biomass. The Chl *a* concentration also showed positive correlations with phosphate and silicate (phosphate: R = 0.73, *p* < 0.01; silicate: R = 0.598, *p* < 0.01). Current velocity had no measurable influence on community variation in different waters (*p* > 0.3). Eukaryotic as well as *Imitervirales* communities in the AS and NECS were well separated from each other in a similar way as geographic distribution and TS diagram showed (Fig. S2).

**Fig. 4.**
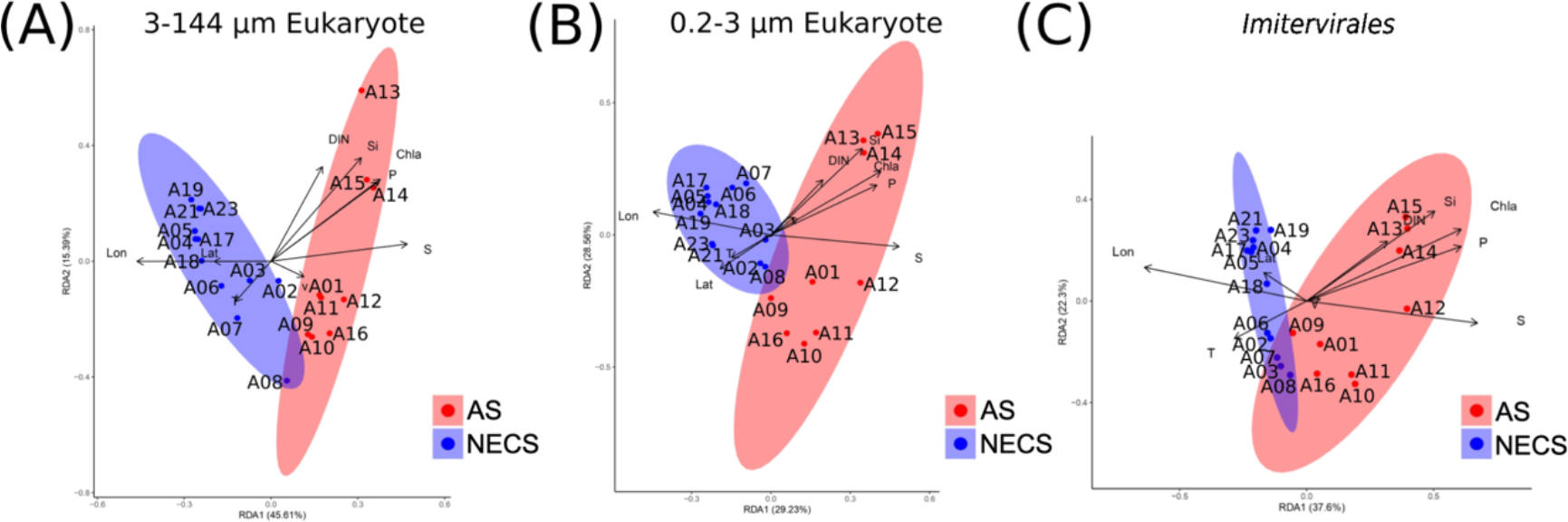
dbRDA (distance-based redundancy analysis) ordination diagram of (A) 3-144 µm eukaryotic community based on 18S ASVs; (B) 0.2-3 µm eukaryotic community based on 18S ASVs; (C) *Imitervirales* community based on *polB* gene ASVs. Abbreviation of water types: NECS: northeastern Chukchi Sea; AS: adjacent sea outside Beaufort Gyre. Abbreviation of geographic and environmental factors: Lat: latitude; Lon: Longitude; T: temperature; S: salinity; DIN: dissolved inorganic nitrogen (sum of ammonia), nitrite and nitrate; P: phosphate; Si: silicate; Chl *a*: chlorophyll *a*; v: density current velocity.

According to the Mantel and partial Mantel tests, eukaryotic communities in both the large (3-144 µm) and small (0.2–3 µm) size fractions correlated significantly with *Imitervirales* communities in both the NECS and AS sites, even when the potential effects of spatial and environmental autocorrelations were removed (q < 0.05) (Table 1). Geographical distance was also a significant factor explaining the eukaryotic communities in the small fraction (q < 0.05), although no significant correlation was found for the large size fraction. For both the size fractions, environmental factors were significant explanatory variables for the eukaryotic communities among the AS sites, whereas no correlation was detected between environmental factors and eukaryotic communities in the NECS sites. The Mantel test was also performed on the eukaryotic 18S communities and each environmental factor (Table S7 and S8). All the environmental factors in the NECS sites were not significantly correlated with the eukaryotic 18S communities in the two size-fractions. In the AS sites, only phosphate and silicate were significantly correlated with the eukaryotic 18S communities.

**Table 1.** R value of Mantel test among different eukaryotic 18S communities in two types of water (separated based on temperature-salinity diagram), geographic factors, environmental factors (including temperature, salinity, dissolved inorganic nitrogen (NO_2_+NO_3_+NH_4_), phosphate and silicate) and *Imitervirales polB* gene community. “*Imitervirales*/geographic” and “*Imitervirales*/environmental” represent the R value of Partial Mantel test, influences of geographic or environmental factors were removed. (q-values were listed in Supplement Table S6)

|  | NECS (N=12) | AS (N=9) |
| --- | --- | --- |
| **3-144μm eukaryote** |  |  |
| geographic factor | 0.30 | 0.34 |
| environmental factor | -0.02 | 0.75** |
| *Imitervirales* | 0.66** | 0.70** |
| *Imitervirales*/geographic | 0.63** | 0.66** |
| *Imitervirales*/environmental | 0.66** | 0.50* |
| **0.2-3μm eukaryote** |  |  |
| geographic factor | 0.44** | 0.30* |
| environmental factor | -0.06 | 0.86** |
| *Imitervirales* | 0.65*** | 0.87*** |
| *Imitervirales*/geographic | 0.61*** | 0.86*** |
| *Imitervirales*/environmental | 0.65*** | 0.90*** |

\*: q<0.05; \*\*: q<0.01; \*\*\*: q<0.001.

## Discussion

### Basic environmental parameters and phytoplankton biomass

Oligotrophy is one of the common features of surface sea water in the NECS. Annual data (2008 to 2010) near the NECS indicated that concentration of nitrate plus nitrite in surface sea water of the study area were mostly depleted in the summer with the values between 0.01 and 0.1 µmol·L^−1^ (Fujiwara et al., 2014). In our study, concentrations of nitrate plus nitrite also showed low values (≤ 0.14 µmol·L^−1^) except for some bloom samples in the AS (0.27–3.12 µmol·L^−1^) (Supplement Table S1). The Chl *a* concentration, which is a proxy of phytoplankton biomass, was also low at the nutrient-depleted stations (0.02–1.70 mg·m^-3^) (Supplement Table S1), suggesting the growth of phytoplankton was limited by nutrient availability (Ko et al., 2020). These values were also consistent with recent Chl *a* data at the corresponding area obtained from satellite (< 0.4 mg·m^-3^) (Lee et al., 2019).

We separated the seawater sampling sites into two groups, NECS and AS, based on the geographical locations. The grouping was also supported by the TS diagram in which the NECS sites were characterized by lower salinity (Fig. S2). The Beaufort Gyre, which influences water properties in the NECS, is the greatest freshwater reservoir in the Arctic (Proshutinsky et al., 2019). On the other hands, samples having higher salinity were classified into the AS, because these locations would be more influenced by oceanic water masses and current regimes including the Pacific Water from the south and Atlantic Water from the west (Jones, 2001; Woodgate, 2013). It is also showed that the main reason of an increase of salinity and nutrient concentrations (resulting from summer algal blooms) in the oligotrophic northeastern Chukchi Sea surface water should be the intrusion of Atlantic cold saline water (Jung et al., 2021).

### Community structures of microbial eukaryotes and *Imitervirales*

The eukaryotic communities were generally dominated by dinoflagellates, diatoms and other alveolates in the large size fraction (3-144 µm) and by ciliates and chlorophytes in the small size fraction (0.2–3 µm) (Fig. 3A and B). The dominance of these groups was roughly consistent with the previous studies that examined the microbial eukaryotic community structures in the Arctic Ocean by the molecular techniques (Comeau et al., 2011; Lovejoy et al., 2006; Marquardt et al., 2016; Onda et al., 2017; Xu et al., 2020) and the satellite ocean color remote sensing (Fujiwara et al., 2014; Lee et al., 2019). Diagnostic pigment signatures have indicated that prasinophytes (Chlorophyta) were the dominant phytoplankton group in the northern Chukchi Sea, while diatoms and dinoflagellates were dominant in the southern Chukchi Sea (Fujiwara et al., 2014). Diatoms and chlorophytes are the common components of spring bloom in the Arctic Ocean (Von Quillfeldt, 2000). In our study, the phytoplankton communities in the bloom sites were dominated by unclassified marine alveolates (45.9% relative abundance) and diatoms (13.5%) in larger size fraction (Fig. 3A). The representative sequence of the unclassified marine alveolate ASV was best hit to the dinoflagellate *Heterocapsa rotundata* in the NCBI Reference RNA sequences database (2021/7/7 updated) (100% sequence similarity). Although this dinoflagellate species has been detected typically in the temperate estuaries (Kyeong et al., 2006; Millette et al., 2015), it was also found to be common near the study area by using microscopic technique (Ardyna et al., 2017).

Unanticipatedly high proportion of metazoan sequences were found in 3-144 µm eukaryotic group (Fig. S3). Most of them belong to copepods, which are predominant zooplankton in the Arctic Ocean (Kosobokova et al., 2011; Wang et al., 2019). However, body sizes of adult free-living copepods are usually above 200 µm, which cannot pass through the pre-filtration mesh. Although some of the copepod species (e.g., *Sphaeronellopsis monothrix*, 110 µm) are even smaller, they are the parasite of marine ostracods (Bowman & Kornicker, 1967). It is reported that smaller eggs of copepods are produced by the adults in spring and summer, and some of these may be float to the surface layer (Hirche & Niehoff, 1996). Thus, one possible explanation for the dominance of metazoan sequences is the emergence of the larvae/eggs in the seawater.

We detected significant differences in the eukaryotic community between the NECS and AS for both size fractions by the dbRDA analysis (ANOSIM, *p* < 0.01) (Fig. 4A and B). In the large size fraction, communities of the NECS sites were consistently dominated by dinoflagellates, whereas the relative abundance of dinoflagellates tended to be lower at the AS sites (Fig. 3A). In the small size fraction (Fig. 3B), community difference between two sampling regimes was not obvious at the phylum level, despite a clear separation by the dbRDA (Fig. 4B). These results suggest that distinct ecosystem structures between the NECS and AS is likely caused by the current systems and associated physicochemical characteristics. We also evaluated eukaryotic communities from the two melt ponds on sea ices, which were located nearby the two northernmost seawater sites (Fig. 3C). The communities were largely distinct from the seawater communities, most likely reflecting the difference in salinity between freshwater and seawater (Xu et al., 2020).

Besides eukaryotes, *Imitervirales* communities were analyzed in our study (Fig. 3C). Among *Imitervirales*, clades 2, 6, and 7 were abundant lineages at most of the sampling sites (Figs 3C and S5). Intriguingly, these three dominant clades do not include any reference species of *Imitervirales*. A previous study reported that the Arctic Ocean is a hot spot of the endemic NCLDVs including *Imitervirales* (Endo et al., 2020); the dominant phylotypes detected in our study may support the high uniqueness of *Imitervirales* phylotypes in the study area. It is suggested that the geographical distribution of viruses follow those of the host species (Ibarbalz et al., 2019), the endemic feature is partly derived from the uniqueness of host eukaryotic species. Community compositions of *Imitervirales* were also differentiated between the NECS and AS stations by dbRDA analysis, as with the eukaryotic communities (Fig. 4C). Expectedly, NMDS analysis (Fig. S6) clearly separated the *Imitervirales* communities in the Arctic sites from those collected from coastal seawater and a hot spring in Japan, which were evaluated using the same MEGAPRIMER method. This separation would be due to the difference in host communities which are primarily determined by the environmental conditions.

### Loose association between environmental variables and eukaryotic community

In this study, salinity was the primary factor used for dividing the sites between the NECS and AS (Fig. S2). Eukaryotic communities were also clearly separated between the NECS and AS (Fig. 4A and B), indicating that the compositions of eukaryotes were strongly influenced by the physical factors in the study area. Thus, we separately assessed the relationship between eukaryotic community and environmental variables or *Imitervirales* community for the NECS and AS to eliminate possible autocorrelation caused by the difference of eukaryotic communities among different water regimes.

In the AS sites, eukaryotic community was strongly correlated with environmental factor, but less correlated with geographical distance (Table 1). This suggests that the community was more affected by physicochemical environmental properties rather than dispersal events such as lateral advection among these sites. In fact, only the phosphate and silicate were significantly correlated with eukaryotic communities in the AS sites (Table S7 and S8). On the other hand, environmental factors (Table 1, S6, S7 and S8) did not show any association with eukaryotic communities in the NECS sites, whereas the effect of geographical distance was comparable to that detected in the AS sites. This indicates that other factors may be more important in making up the eukaryotic communities in the Beaufort Sea basin. In our study, all the sampling sites in the NECS were oligotrophic, and in some locations, the concentrations of nutrients were below the detection limit. Additionally, although temperature and salinity tend to be the key factors for microbial eukaryotic community structure and distribution in marine ecosystem (Caron et al., 2016; Sherr et al., 2007), these variables did not largely vary among the NECS sites. The low variation in environmental condition may cause the lack of correlation between environmental variables and the eukaryotic community.

### Tight association between *Imitervirales* and the microbial eukaryotic community

In contrast to environmental variables, *Imitervirales* communities were consistently correlated with eukaryotic communities in both the NECS and AS regions (Table 1). Notably, the correlation coefficients were rarely influenced by the geographical and environmental factors, suggesting that *Imitervirales* were associated with the eukaryotes in both types of water independently from environmental factors. This trend was most pronounced at the stations in the NECS, where environmental variables were relatively stable and had no correlation with eukaryotic community variations. Our results support the idea that the communities of *Imitervirales* and eukaryotes are actively interacting and co-varying without detectable influence from the environmental conditions even in oligotrophic and homogeneous environments.

It has been suggested that biological interactions, such as predator-prey and symbiotic interactions, are responsible to determine community structure and the dynamics of microbes (Chaffron et al., 2020; Lima-Mendez et al., 2015). Additionally, viruses have been proposed as a key factor influencing the protist communities as they can impose top-down controls on their specific host populations (Brussaard et al., 1996; Nagasaki et al., 1994). Recent studies using Mantel statistics or co-occurrence network analysis indicated that *Imitervirales* are tightly associated with a variety of protist lineages at a global level (Endo et al., 2020; Meng et al., 2021), although only little of them have been isolated (Mihara et al., 2018). Our 18S rDNA barcoding revealed that chlorophytes and haptophytes, both of which are known host lineages of *Imitervirales*, were major protists in the small size fraction. Although the dominating clades in the large-sized eukaryotic communities such as dinoflagellates, ciliophora, and diatoms have not yet been reported as host lineages, these groups were predicted to be the most closely linked host group for *Imitervirales* from a global scale network analysis (Meng et al., 2021). Considering the highest proportion of *Imitervirales* among NCLDVs in the global ocean and their potential role as a top-down factor on host populations, relative compositions of the host lineages may well result from the combination of a variety of specific infections of NCLDVs and other viruses.

In the Arctic Ocean, an increase in sea surface temperature and decrease in sea ice cover are progressing (Peng et al., 2020; Praetorius et al., 2018). These climate change has been shown to be associated with the shift of eukaryotic community structure as well as the increase of biomass and the potential loss of biodiversity in the past decade (Arrigo & van Dijken, 2015; Li et al., 2009; Majaneva et al., 2012), although another study suggests a decreasing tendency on biomass (Hill et al., 2013). Increased temperature may provide competitive advantage to small nanophytoplankton over larger phytoplankton, resulted in an increase of the contribution of small phytoplankton in the community (Hare et al., 2007; Li et al., 2009). Our study showed that the association with *Imitervirales* community was generally higher for the small-sized plankton community than for the large-sized community, implying the role of *Imitervirales* in structuring the eukaryotic community in the study area may become increasingly important in a future. However, it should be noted that virus-host interactions can be influenced by the environments, especially temperature (Demory et al., 2017, 2021).

## Experimental procedures

### Sampling sites and processes

During the Arctic Ocean Cruise of the IBRV Araon 2018 of Korean Polar Research Institute (KOPRI), surface water samples were collected (SBE32 carousel water sampler) at 21 stations from 6th to 22nd of August 2018. Environmental parameters including salinity, temperature, Chl *a* and nutrient concentration were obtained in parallel. Salinity and temperature were measured by the CTD sensors in situ measurement of seawater. For Chl *a*, seawater samples were collected in the upper 100 m depth and filtered through 47 mm GF/F filters, then was extracted with 90% acetone (Jung et al., 2021). Chl *a* was measured by fluorometer (Trilogy, Turner Designs, USA) (Lee et al., 2016). For nutrient concentration measurement, 50 ml seawater sample for each site was collected by conical tube, stored at 4°C. Nitrite, nitrate, ammonia, phosphate, and silicate were measured using a four-channel continuous auto-analyzer (QuAAtro, Seal Analytical) followed the Joint Global Ocean Flux Study (JGOFS) protocols (Gordon et al., 1993). Nutrient concentrations under detection limit and lower than 0.005 µmol·L^−1^ were considered 0.

Seawater (1 L) for the DNA analysis was collected from 2 m depth with Niskin-bottles attached to a CTD–CMS system for all stations except at two closed melt ponds (500 mL), where water samples were collected just below the surface by bucket. Collected seawater was prefiltered with a 144 µm pore-size mesh to remove large particles (prewashed with ultrapure water). Two liters of water were separated into two replicates on average, then were filtered through 3 µm Millipore membrane filter by air pump (< 0.03MPa) for larger size fraction, further filtered through 0.2 µm Millipore membrane filter with the same method for smaller sized fraction. The membrane filters were transfer to 1.5 mL microtubes and then stored in −20°C on board and then transferred to the laboratory while continuously kept at −20°C.

### DNA extraction and purification

DNA extraction and purification were performed following (Endo et al., 2013; Endo et al., 2018). Briefly, each membrane filter was thawed at room temperature and was put into the 1.5 mL microtubes with glass beads and XS buffer. The cells on filter were crushed with a beads beater and the mixture was incubated at 70°C for 60 min. Glass beads were removed from mixture after centrifugation. 600 µL isopropanol were added to the supernatant and mixed. The precipitated DNA was purified with NucleoSpin gDNA Clean-up Kit (Macherey-Nagel). Finally, the purified DNA was dissolved in low TE buffer and stored at −20°C.

### Eukaryotic 18S gene amplification and purification

Eukaryotic 18S rRNA gene V4 region fragments were amplified from extracted DNA of both 3 µm and 0.2 µm size fractions using primer E572F (5’-CYGCGGTAATTCCAGCTC-3’) and E1009R (5’-AYGGTATCTRATCRTCTTYG-3’) (Comeau et al., 2011) with attached Illumina MiSeq 300 PE overhang reverse adapters as described in Illumina metagenomic sequencing library preparation protocols.

12.5 µL 2x KAPA HiFi HotStart ReadyMix was mixed with 5 µL 1 µmol·L^−1^ amplicon PCR forward primer, 5 µL 1µmol·L^−1^ amplicon PCR reverse primer and 2.5 µL diluted DNA samples (0.25 ng·µl^-1^), and were added into a PCR tube (final volume 25 µL). The amplification was performed for each sample with the following temperature cycling condition: initial denaturation at 98°C for 30 sec was followed by 30 cycles of denaturation at 98°C for 10 sec, annealing at 55°C for 30 sec and 72°C for 30 sec. A final extension step was at 72°C for 5 min.

Amplicons were purified with magnetic beads (Agencourt AMPure XP beads, Beckman Coulter, Inc.). The purified DNA were dissolved in 25 µL ultrapure water and stored at −20°C.

### *Imitervirales polB* gene amplification and purification

The degenerated 82 *polB* primer pairs (MEGAPRIMER, Supplement Table S3) were used to amplify the *polB* gene of *Imitervirales* from 0.2 µm membrane filter DNA samples (Li et al., 2018). A previously optimized amplification method named “MP10” (Supplement Table S4) was performed. amplification protocol, materials and temperature cycling condition were the same as a previous work (Prodinger et al., 2020).

After amplification, we merged all the eight amplicons generated from the same DNA sample using ethanol precipitation (Prodinger et al., 2020). Finally, the DNA precipitation was air dried for around 10 min and resuspended in 25 µL ultrapure water. Gel extraction was performed to remove unspecific amplification products. Gel electrophoresis was made by 2% agarose gel. The gel was then stained in 5000x diluted SYBR gold buffer for 12 min. 400-500 bp visible bands were cut from the gel. The Promega’s Wizard SV Gel and PCR Clean-Up System was used to perform gel extraction according to the marker’s protocol. DNA was dissolved in 25 µL ultrapure water, stored at −20°C.

### Index PCR, library construction and sequencing

Index PCR was performed following the Illumina Miseq platform protocol. Produced amplicons of 3-144 µm eukaryotes and *Imitervirales* were purified with the magnetic beads (Agencourt AMPure XP beads, Beckman Coulter, Inc.). Finial DNA production was dissolved in 27.5 µL ultrapure water, stored at −20°C less than 24h. Produced amplicons of 0.2-3 µm eukaryotes were purified by gel, performed by Macrogen Corp. Japan.

DNA concentration was measured by Qubit HS (high-sensitive) kit. Library was denatured following the standard MiSeq normalization method provided by Illumina. The MiSeq Reagent Kit v2 and NaOH were used for the library with final DNA concentration of 2 nM. Paired-end sequencing was performed on the MiSeq platform.

### Sequence processing and bioinformatic analysis

Eukaryotic 18S sequences are processed with QIIME2 (version: 2019.10) (Bolyen et al., 2019). 260 bp of left pair reads and 220 bp of right pair reads were trimmed. DADA2 was used to cut primer sequences, merging the paired end reads, performing quality control, dereplication, chimera check, and Amplicon Sequence Variants (ASVs) generation (Callahan et al., 2016; Knight et al., 2018). Singleton ASVs were removed. Taxonomic annotation was done with QIIME2’s vsearch (Rognes et al., 2016) plugin and the SILVA 132 small subunit with 97% similarity database (Quast et al., 2013) at 97% identity for species eukaryotic 18S datasets. Dominant ASVs (reads percentage over 0.50% of each size fraction) were again searched by blastn (Altschul et al., 1990) in NCBI Reference RNA sequences dataset, result include detailed linage information with highest identity value was selected.

For the *Imitervirales* sequences, MAPS2 (*Mimiviridae* Amplicon Processing System) was used for sequence analysis (Prodinger et al., unpublished). DADA2 was used to check and remove megaprimer sequences, merging, quality control, dereplication, chimera check, and non-singleton ASV output. The ASVs were aligned against *Imitervirales polB* amino acid sequence database (Li et al., 2018). Nucleotide sequences were translated into amino acid sequences and then added to reference alignment using mafft (version: 7.453, parameters: --thread −1 –genafpair--maxiterate 1000) (Katoh et al., 2002; Katoh & Standley, 2013). Sequences which were assigned to the *Imitervirales* were saved for the further analysis, while other sequences were removed. Translated ASVs were placed into the reference phylogenetic tree built by the *polB* sequences from *Tara* Ocean dataset (Endo et al., 2020) by pplacer (version: 1.1. alpha19) (Matsen et al., 2010). Thirteen *Imitervirales* clades were manually defined in the tree, and ASVs were assigned to each clade. Phylogenetic tree was edit and output by iTOL v5.7 (Letunic & Bork, 2019).

### Ecological analysis

Community composition was evaluated based on number of reads of each ASV in every sample. ASVs were then subsampled by the rarefy function (“vegan” package) (Oksanen et al., 2018) in R (version 3.6.3). Relative abundance was represented by the rate of each ASV reads percentage in each sample. Shannon diversity index of eukaryotic and *Imitervirales* community was calculated by R (“vegan” package) based on the subsampled ASV table. ANOVA and Tukey post hoc tests were performed by R (“agricolae” package). Composition bar chart and diversity bar chart with error bars of standard deviation were calculated with Microsoft Excel (version 16.41). The map of sampling stations, temperature-salinity (TS) diagram and heatmap of environmental factors and Shannon diversity were generated by Ocean Data View (ODV, version 5.1.5) (Schlitzer, R., Ocean Data View, https://odv.awi.de, 2018). Biological correlation was performed by dbRDA (distance-based redundancy analysis) function (Legendre & Andersson, 1999), using R (“vegan” package) based on Bray-Curtis dissimilarity. For dbRDA ordination, ASV composition was normalized by Hellinger transformation by decostand function. Spearman’s rank correlation was performed by R (cor.test function) and *p* value was also calculated by R (cor.test function). ANOSIM with 9,999 permutation was performed for biological data grouping test. Results of dbRDA were plotted by “ggord” with 95% confidence interval circle contained samples in different water types. Non-metric multidimensional scaling (NMDS) analysis of *Imitervirales* community was performed by R (monoMDS function) based on Bray-Curtis dissimilarity matrix made with the subsampled ASV table.

Mantel test and partial Mantel test (Mantel, 1967; Smouse et al., 1986) based on Pearson correlation coefficient were performed for calculating the correlation among geographic distance, environmental variables (i.e., a distance matrix combining temperature, salinity, dissolved inorganic nitrogen (DIN, nitrate + nitrite + ammonium nitrogen), phosphate, and silicate), eukaryotic community and *Imitervirales* community, using R (“ade4” package) (Bougeard & Dray, 2018) with permutations of 1,000. Geographic distance between each sampling station was calculated with R (“geosphere” package) from latitude and longitude data. Every environmental variable was normalized by log_10_(x+1) function (x: the value of environmental factor). Euclidean distance of environmental factors and Bray-Curtis dissimilarity of subsampled relative abundances of eukaryotes and *Imitervirales* between sampling sites were calculated with R. All *p* values were adjusted by the Holm’s method (Holm, 1979) using R’s p.adjust function.

## Data availability

The raw reads generated in this study were uploaded to SRA (Sequence Read Archive) database on NCBI website. The accession numbers are from SRR12981736 to SRR12981758 under project ID PRJNA674408 (3-144 µm eukaryotic 18S), from SRR12981654 to SRR12981676 under project ID PRJNA674418 (0.2-3 µm eukaryotic 18S) and from SRR12981759 to SRR12981780 under project ID PRJNA674422 (0.2-3 µm *Imitervirales polB*).

## Acknowledgments

We would like to thank colleagues from Korea Polar Research Institute for the help of sampling and physicochemical parameter determination; Tatsuhiro Isozaki, Kento Tominaga and Hiroaki Takebe from Laboratory of Marine Microbiology, Kyoto University, for helping with DNA sequencing and support of experiment. We also thank the captain and crew of the IBRV Araon Cruise for their support during the cruise. This work was supported by JSPS/KAKENHI (Nos. 18H02279 and 19H05667 to H.O., 17H03850 to T.Y. and H.O., and Nos. 19K15895 and 19H04263 to H.E.), and Scientific Research on Innovative Areas from the Ministry of Education, Culture, Science, Sports and Technology (MEXT) of Japan (Nos. 16H06429, 16K21723, and 16H06437 to H.O.). This research was also supported by the project titled ‘Korea-Arctic Ocean Warming and Response of Ecosystem (K-AWARE, KOPRI, 1525011760)’, funded by the Ministry of Oceans and Fisheries, Korea (KHC, JJ, EJY, SHK). Computational work was completed at the SuperComputer System, Institute for Chemical Research, Kyoto University. The authors declare no conflicts of interest.

---

**Fig. S1.**
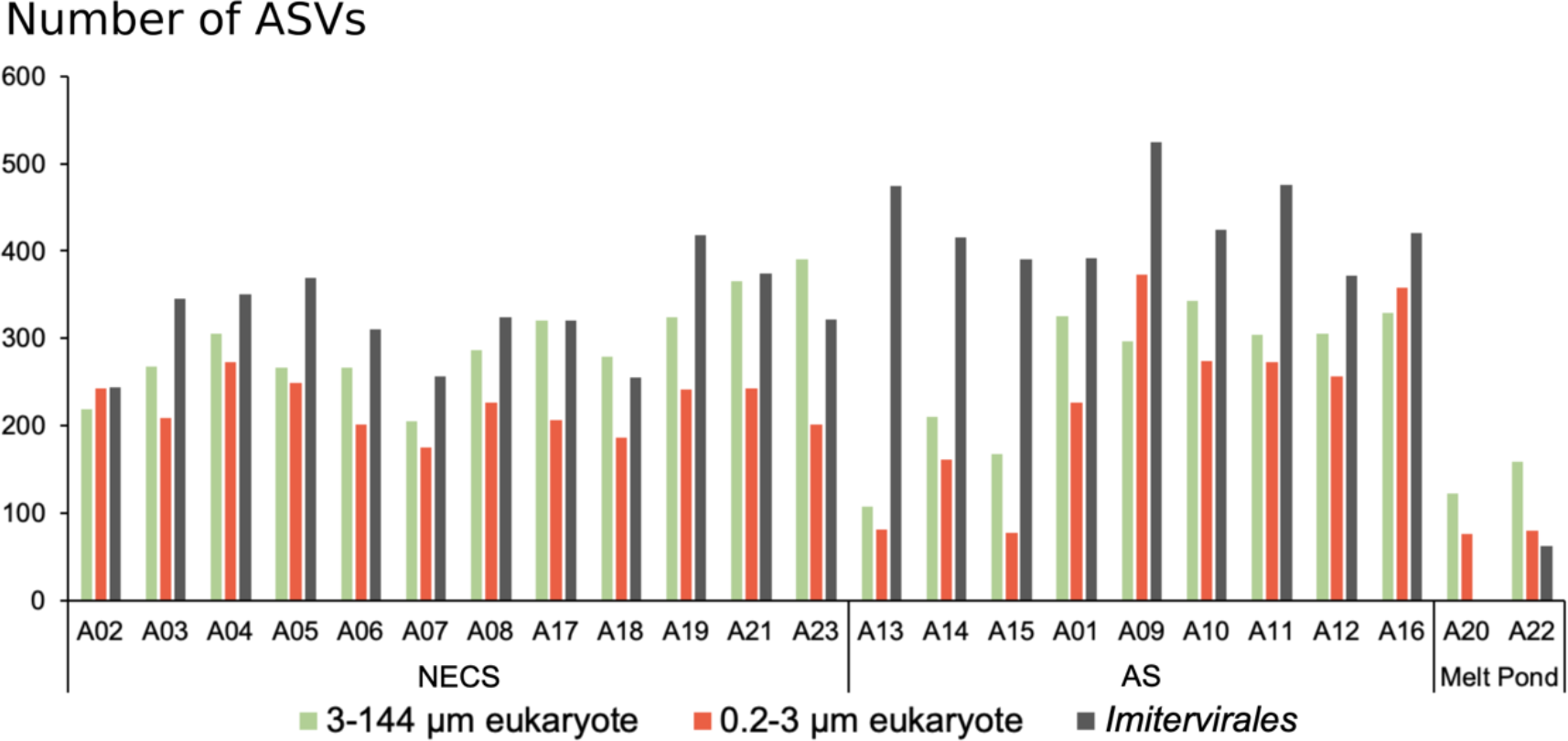
Number of non-singleton ASVs of each sample before subsampling. NECS: northeastern Chukchi Sea; AS: adjacent sea outside Beaufort Gyre.

**Fig. S2.**
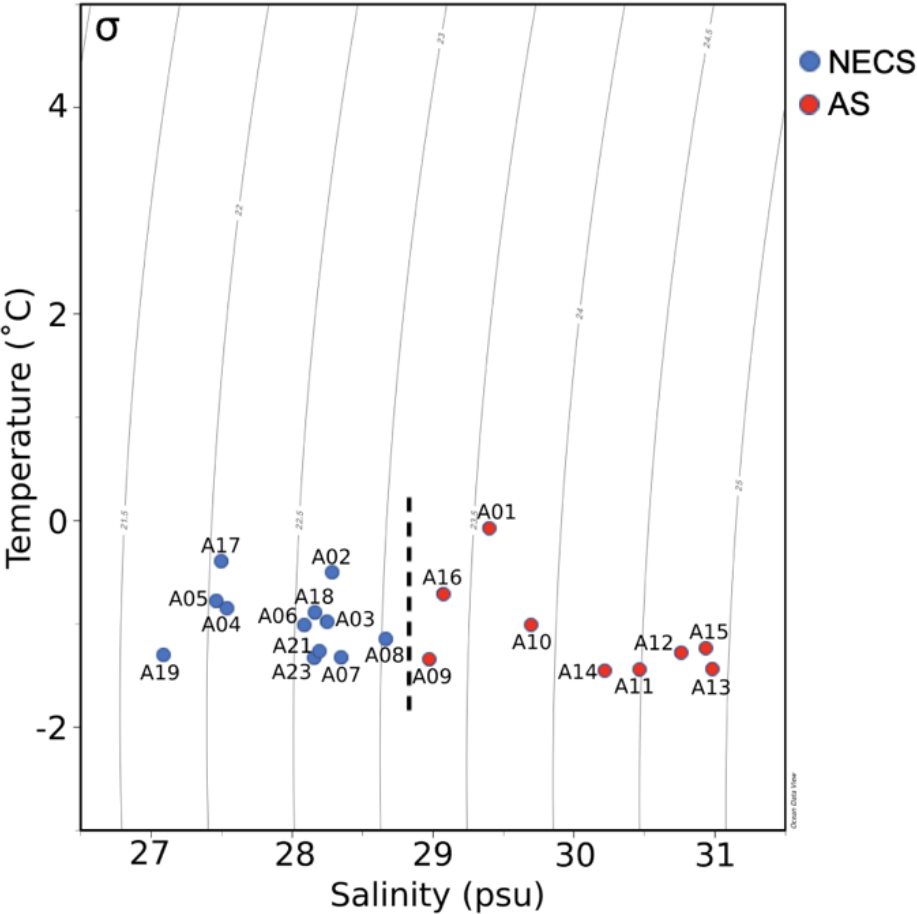
TS diagram of all the sampling sites in this study. T: temperature; S: salinity; σ: density. NECS: northeastern Chukchi Sea; AS: adjacent sea outside Beaufort Gyre. Dashed line separates the different water types.

**Fig. S3.**
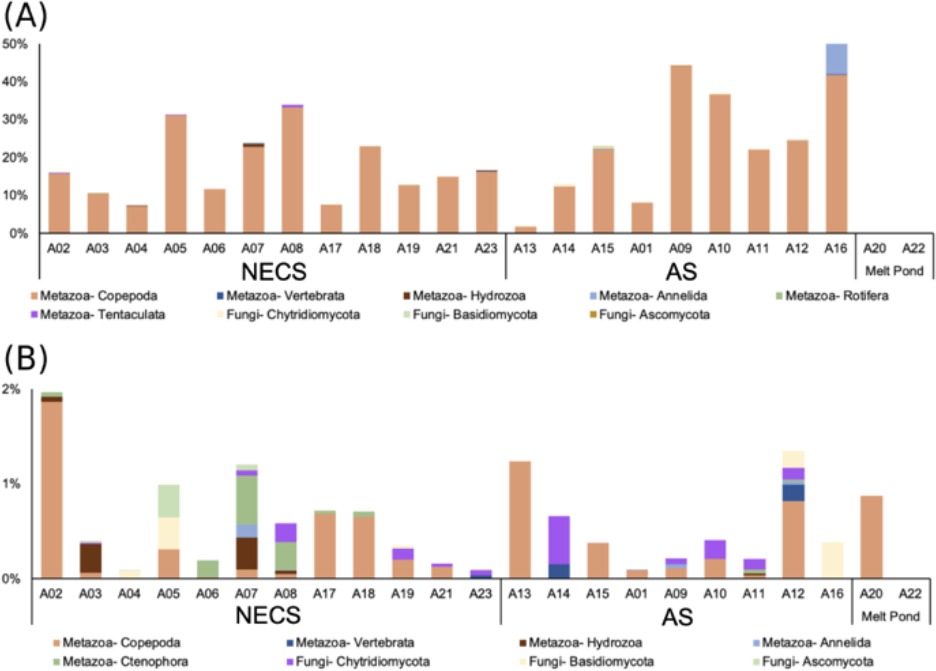
Contribution of metazoa and fungi to the total eukaryotic communities in the (A) 3-144 µm fraction and (B) 0.2-3 µm fraction.

**Fig. S4.**
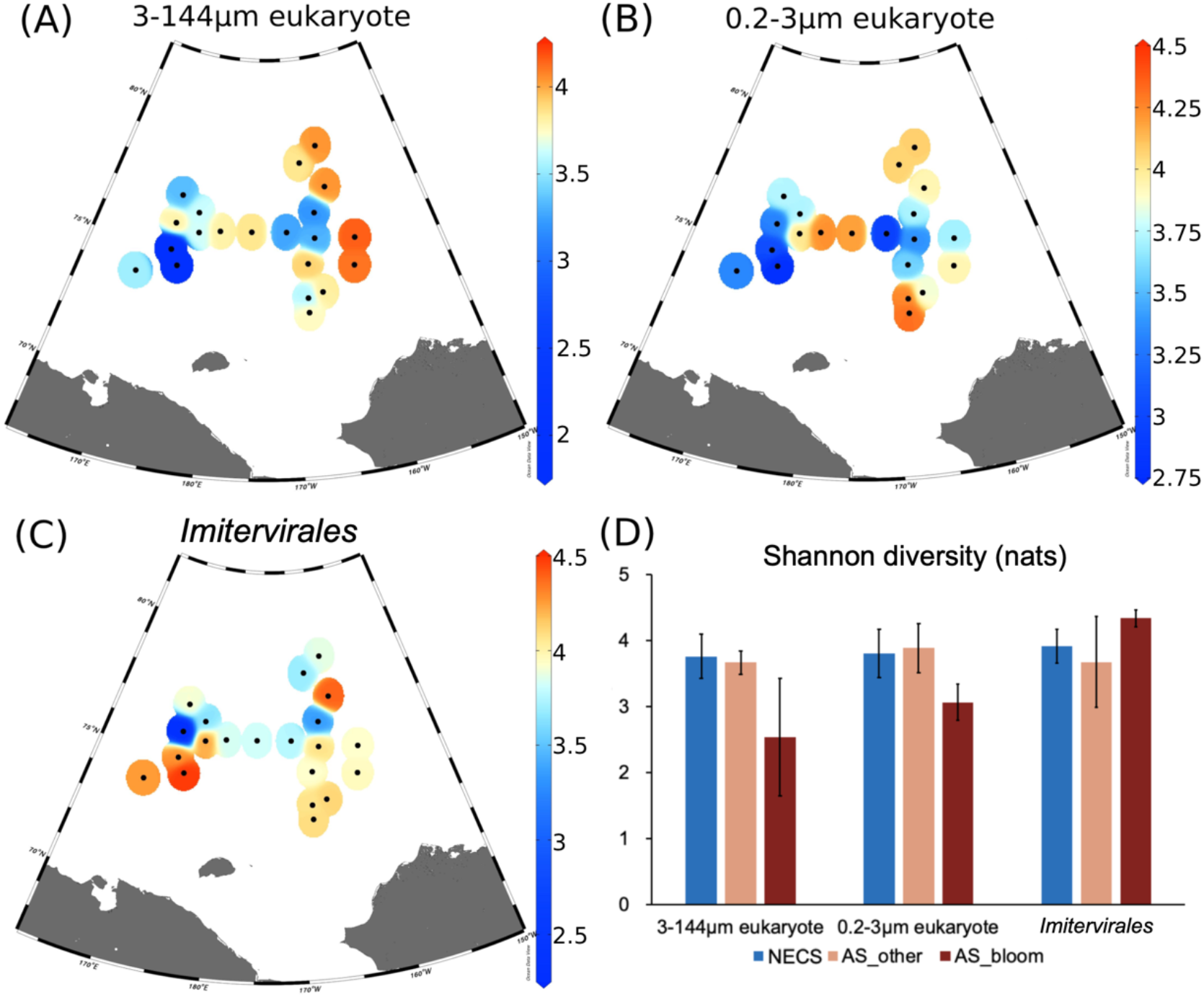
Distribution of Shannon diversity index across sampling sites: (A) eukaryotes in the 3-144 µm fraction; (B) eukaryotes in the 0.2-3 µm fraction; (C) *Imitervirales*. (D) Bar plots summarizing the Shannon diversity in water type for the three communities (error bar of ± one standard deviation). Abbreviation of water types: NECS: northeastern Chukchi Sea; AS: adjacent sea outside Beaufort Gyre.

**Fig. S5.**
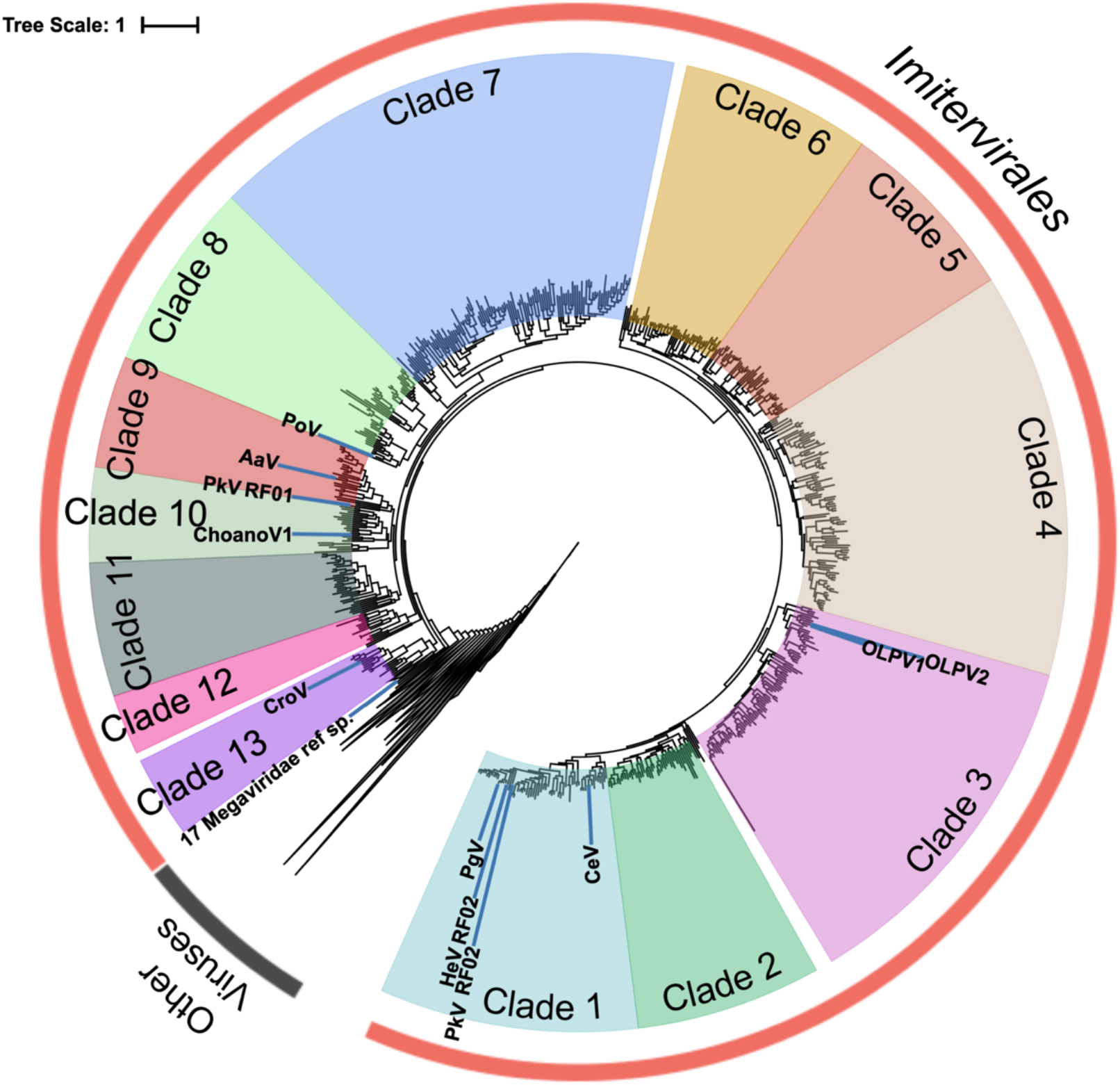
Phylogenetic tree built based on marine virus *polB* gene from *Tara* Ocean dataset. Clade 1 to 13 are belonging to *Imitervirales*. Reference sequence in the thirteen clades: PgV (clade 1) - *Phaeocystis globosa* virus; HeV (clade 1)- *Haptolina ericina* virus; PkV (clade 1 and clade 9) - *Prymnesium kappa* viruses; CeV (clade 1) - *Chrysochromulina ericina* virus; OLPV (clade 3) - Organic Lake Phycodnaviruses; PoV (clade 8) - *Pyramimonas orientalis* virus; AaV (clade 9) - *Aureococcus anophagefferens* virus; ChoanoV1 (clade 10) – Choanovirus1; CroV (clade 13) - *Cafeteria roenbergensis* virus.

**Fig. S6.**
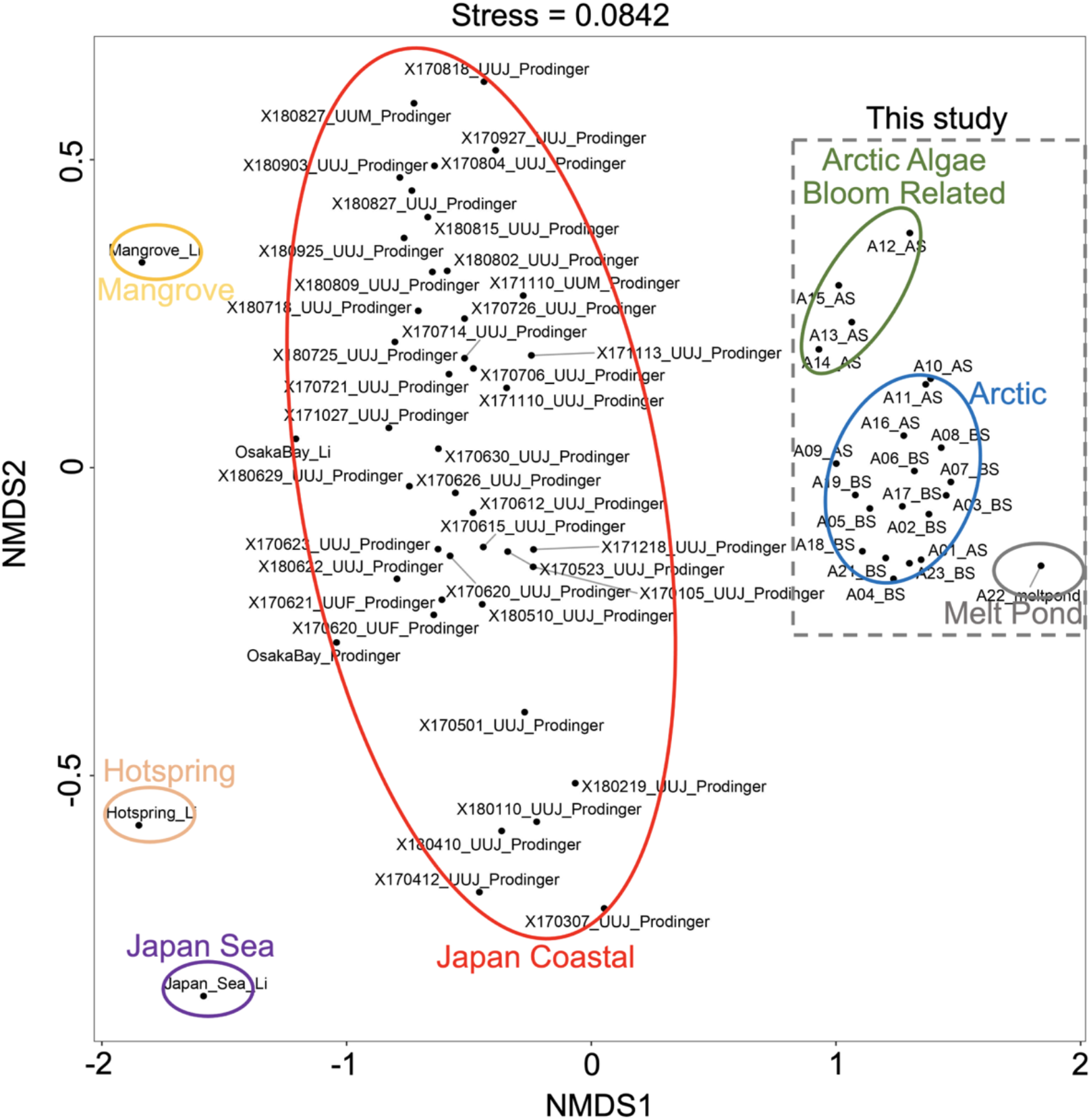
Non-metric multidimensional scaling (NMDS) plot with *Imitervirales* ASVs of Arctic samples in this study, hotspring samples, subtropical coastal sea water samples (Li et al., 2019; Prodinger et al., 2020, Prodinger et al., unpublished). ASVs were firstly subsampled by the minimum number of each sample, and relative abundance was calculated by percentage of ASVs in each sample. Bray-Curtis dissimilarity was used to calculate the distance matrix.

**Fig. S7.**
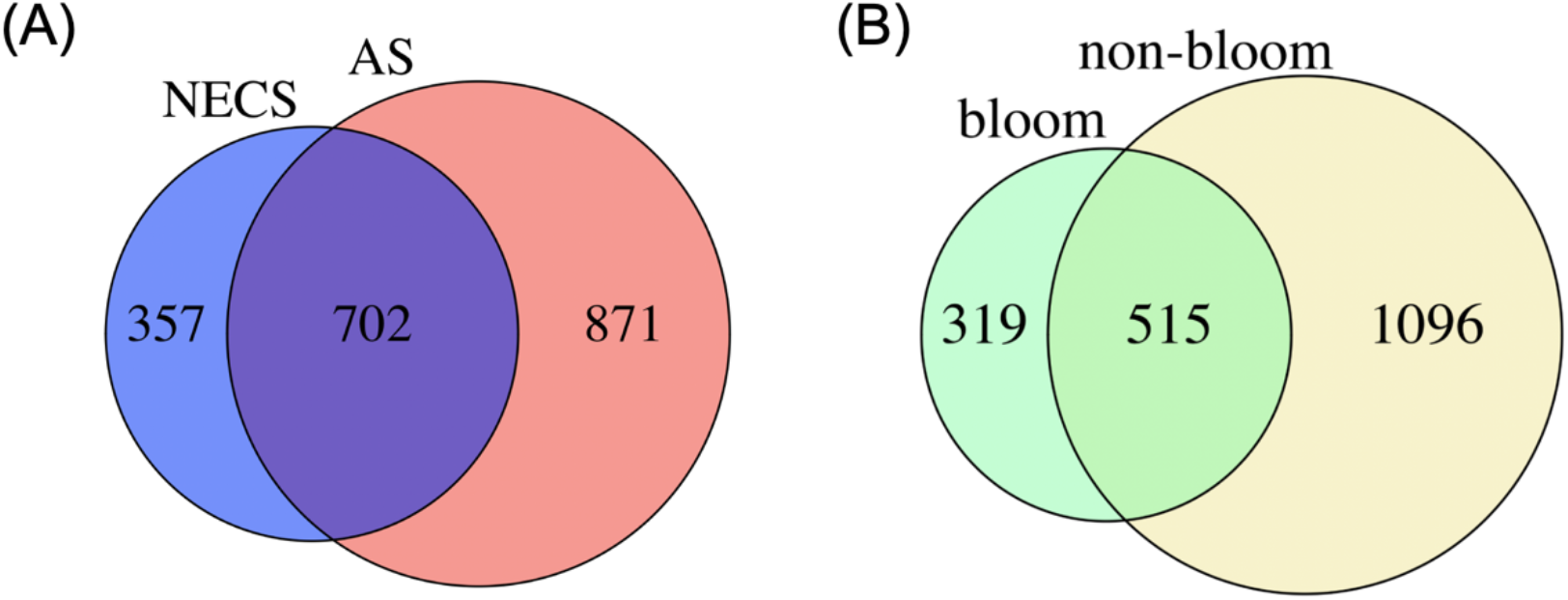
Number of unique and common non-singleton *Imitervirales* ASVs between (A) NECS and AS and between (B) bloom and non-bloom sites.

Dataset S1. Microsoft Excel file includes 10 supplementary tables of sampling data, sequencing data, primer data and secondary result of analysis.

